# *Xanthomonas sontii* sp. nov., a non-pathogenic bacterium isolated from healthy basmati rice (*Oryza sativa*) seeds from India

**DOI:** 10.1101/738047

**Authors:** Kanika Bansal, Amandeep Kaur, Samriti Midha, Sanjeet Kumar, Suresh Korpole, Prabhu B. Patil

**Author notes:** Address correspondence to Prabhu B. Patil. Institute of Infection and Global Health, University of Liverpool, Liverpool, UK. Department of Archaeogenetics, Max Planck Institute for the Science of Human History, Jena, Germany (Sanjeet Kumar). Equal contribution (Kanika Bansal, Amandeep Kaur). **Data submission:** Whole genome sequences of PPL1, PPL2 and PPL3 strains are submitted to NCBI with accession numbers NQYO, NQYP, NMPO respectively.

## Abstract

Three yellow pigmented, Gram negative, aerobic, rod shaped, motile bacterial strains designated as PPL1, PPL2 and PPL3 were isolated from healthy basmati rice seeds. Phenotypic, biochemical and 16S rRNA gene sequence analysis assigned these strains to the genus *Xanthomonas*. The 16S rRNA gene sequence was having 99.59% similarity with *X. sacchari* CFBP4641^T^. However, whole genome based phylogenomic analysis revealed that these strains formed a distinct monophyletic clade with *X. sacchari* CFBP4641^T^ as their closest neighbour. Taxonogenomic studies based on average nucleotide identity (orthoANI) and digital DNA-DNA hybridization (dDDH) values of these strains with type strains (or representative strains) of different *Xanthomonas* species including *X. sacchari* showed below recommended threshold values of ANI (<96%) and dDDH (70%) for species delineation. Furthermore, at the whole genome level, PPL1 and PPL2 were found to be clonal, while PPL3 was not a clonal, but belonging to the same species. Our *in planta* pathogenicity studies revealed that the strains PPL1, PPL2 and PPL3 are non-pathogenic to rice plants. Hence, based on the present study, they form a novel lineage and species associated with rice seeds for which the name *Xanthomonas sontii* sp. nov. is proposed. The type strain for the *X. sontii* sp. nov. is PPL1^T^ (CFBP8688^T^ = ICMP23426^T^ = MTCC12491^T^) and strains PPL2 (CFBP8689 = ICMP23427 = MTCC12492) and PPL3 (CFBP8690 =ICMP23428 = MTCC12492) as other strains of the species.

## Introduction

*Xanthomonas* is a Gram-negative, yellow-pigmented plant associated bacterium that infects economically important crops such as rice [1], pomegranate [2], citrus [3], pepper, cabbage [4], banana [5] etc. It is a complex genus comprising of 33 species (http://www.bacterio.net/), which are further classified into 150 pathovars [6]. In 1997, Vauterin *et al*., divided these *Xanthomonas* species into three clusters based on 16S rRNA gene sequence phylogenetic analysis [7]. Besides main core *Xanthomonas* cluster, *X. albilineans, X. hyacinthi, X. theicola*, and *X. translucens* grouped as second cluster, whereas *X. sacchari* formed a distinct phylogenetic cluster. However, advent of next generation sequencing technology and introduction of robust whole genome based tools like orthologous average nucleotide identity (orthoANI) and digital DNA-DNA hybridisation (dDDH) have revolutionized the field of bacterial taxonomy [8-10]. Infact, these methods are refining comprehensive taxonomic framework, which are essential for developing diagnostic strategies and understanding host-pathogen relationships in management of crops [8].

*Xanthomonas* is emerging as a serious threat for economically important crops. *X. oryzae* pv. oryzae, *X. campestris, X. axonopodis* pv. manihotis were considered in the top 10 plant pathogenic bacteria [11]. Among these, *X. oryzae* causes bacterial blight disease to rice plants resulting in 30-50% decrease in rice yield every year [12], [13], [14]. Other than pathogenic *Xanthomonas* strains that cause disease in rice plants, some of the non-pathogenic *Xanthomonas* strains have also been identified from rice plants [15], [16]. Most of these non-pathogenic strains reported were largely characterized based on phenotypic and biochemical analysis providing limited information.

In the present study, we report isolation and characterization of three creamish-yellow pigmented bacterial strains from healthy basmati rice (*Oryza sativa*) seeds. Genome based polyphasic analysis supported with pathogenicity tests revealed that these strains are non-pathogenic to rice and belong to a novel species, for which we propose *Xanthomonas sontii* sp. nov. These non-pathogenic stains are widely over-looked due to their less economic importance. However, these non-pathogenic isolates were isolated from rice plant, where their pathogenic counterparts (*X. oryzae*) causes infection and are devastating worldwide. Hence, identification and detailed analysis of these non-pathogenic strains can provide important insights into the lifestyle adapted by these strains and virulence mechanisms of their pathogenic counterparts.

## Materials and methods

### Bacterial strain isolation and culture conditions

Strains were isolated from surface sterilized healthy rice seeds (Pusa basmati 1121 variety) that were collected from Fazilka, Punjab, India (30.4036° N, 74.0280° E). For bacterial strain isolation, surface sterilized seeds were partially crushed in 0.85 % NaCl (normal saline) using sterile mortar and pestle. The mixture was then suspended in 50 ml of solution [17]. The solution was incubated for 2 h at 28°C and serial dilutions performed up to 10^-6^ and different dilutions (100µl) were plated on media like nutrient agar (NA), peptone sucrose agar (PSA) and glucose yeast extract calcium carbonate agar (GYCA). Plates were incubated at 28°C up to 6 days. Bacterial colonies isolated were grown and maintained on PSA medium.

### Phenotypic and biochemical characterisation

We analysed the morphology of strains by observing presence of flagella using transmission electron microscopy. PPL1 strain was grown in nutrient broth and incubated at 28 °C for 20 h. Subsequently, cells were harvested by centrifugation at 2000 rpm for 10 minutes. Cell pellet was washed twice with phosphate buffer saline (1X PBS) and finally resuspended in PBS. Bacterial suspension was placed on carbon-coated copper grid (300 mesh, Nisshin EM Co., Ltd.) for 15 minutes. The grid was then negatively stained for 30 seconds with 2% phosphotungstic acid, dried and examined under JEM 2100 transmission electron microscope (JEOL, Tokyo, Japan) operating at 200 kV.

Further, biochemical characterization such as carbohydrate utilization, acid production and various enzymatic activities were performed using OMNILOG GEN III system (BIOLOG) according to manufacturer’s instructions.

### DNA extraction, genome sequencing, assembly and annotation

Genomic DNA extraction was carried out using ZR Fungal/Bacterial DNA MiniPrep kit (Zymo Research, Irvine, CA, USA). Qualitative assessment of DNA was performed using NanoDrop 1000 (Thermo Fisher Scientific, Wilmington, DE, USA) and agarose gel electrophoresis. Quantitative test was performed using Qubit 2.0 fluorometer (Life Technologies). Nextera XT sample preparation kits (Illumina, Inc., San Diego, CA, USA) were used to prepare Illumina paired-end sequencing libraries (250 x 2 read length) with dual indexing adapters. In-house sequencing of the Illumina libraries was carried out on Illumina MiSeq platform (Illumina, Inc., San Diego, CA, USA). Adapter trimming was performed automatically by MiSeq control software (MCS), and remaining adapters were detected by NCBI server and were removed by manual trimming. Sequencing reads were *de novo* assembled into high quality draft genome on CLC Genomics Workbench v7.5 (CLC bio, Aarhus, Denmark) using default settings. Genome annotation was performed by NCBI PGAP pipeline (http://www.ncbi.nlm.nih.gov/genome/annotation_prok).

### Phylogenomic and taxonogenomic analysis

The 16S rRNA sequence of the strains was fetched from genome sequence and its comparision with validly published reference bacteria was carried out using Eztaxon (https://www.ezbiocloud.net/). Core genome tree was constructed using PhyML[18]. Core genome alignment was obtained using Roary v3.11.2 [19] with identity cutoff 60%. The core gene alignment was converted into phylip format using SeaView v4.4.2-1 [20] and then, newick tree was obtained using PhyML. *Stenotrophomonas maltophilia* ATCC13637 was used as an outgroup. Taxonogenomic analysis of all type strains or representative strains of *Xanthomonas* was performed using OrthoANI v1.2 [21] values calculated by using USEARCH v5.2.32 [22] and dDDH were calculated using Web tool GGDC 2.0 (http://ggdc.dsmz.de/distcalc2.php)

### *In planta* pathogenicity test

PPL1, PPL2, PPL3 and BXO1 were grown to saturation in PSA media and inoculated on 30 days old rice plant (PUSA-basmati 1121) variety. Inoculation was performed by dipping scissors in bacterial culture and clipping tips of rice leaves. After 14 days, infection was assessed by measuring length of lesions on leaves. Here, BXO1 was positive control and PBS was used as negative control. Pathogenicity data of each isolate was obtained from 10 inoculated leaves based on two independent experiments.

## Results and discussion

### Phenotypic and biochemical characterisation of PPL1, PPL2 and PPL3

All strains PPL1, PPL2 and PPL3 were isolated form glucose yeast extract calcium carbonate agar (GYCA) media after 24 h of incubation at 28 °C. Colonies appeared as yellow, round, smooth, convex and circular. All strains were Gram-negative, rod shaped bacteria with monopolar flagella, as shown in figure 1.

**Figure 1:**
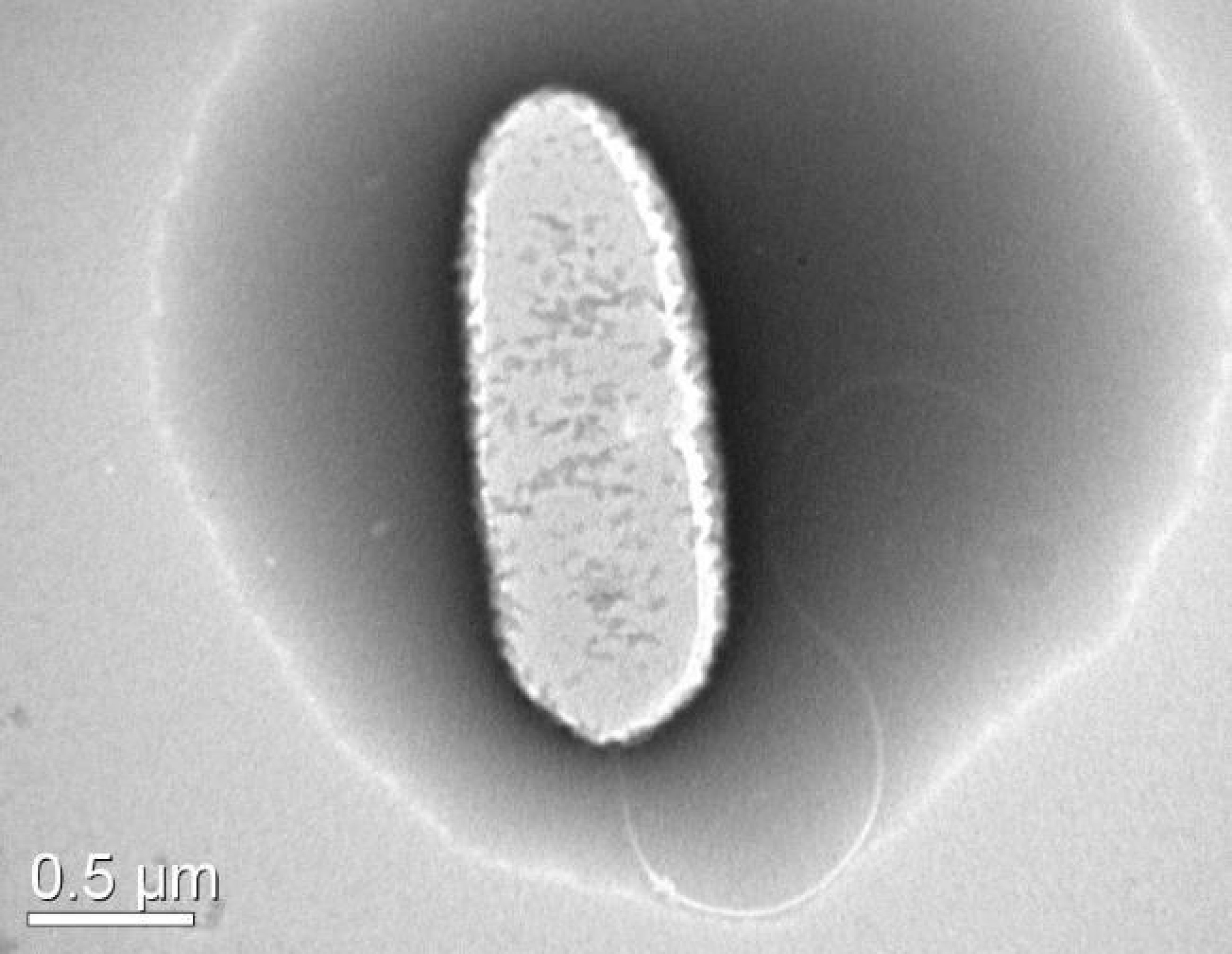
Transmission electron microscopy image of PPL1^T^ strain with monopolar flagella.

Major biochemical characteristics of PPL1, PPL2 and PPL3 strains along with their closest neighbour *X. sacchari* NCPPB 4341^T^ were determined using BIOLOG GEN III MICROPLATE™ and compared with *X. albiliniens* LMG 494^T^ [23] (table 1). All three strains grew well between 20°C to 37 °C with optimum temperature 28 °C. No growth observed at 50 °C. Further, strains were able to grow at pH 6.0 and up to 4% NaCl whereas no growth observed at pH 5.0 and 8% NaCl. In BIOLOG results, all strains PPL1, PPL2 and PPL3 were positive for utilization of D-maltose, D-trehalose, D-cellobiose, gentiobiose, sucrose, D-turanose, α-D-lactose, D-melibiose, β-methyl-D-glucoside, D-salicin, N-acetyl-D-glucosamine, α-D-glucose, D-mannose, D-fructose, D-galactose, L-fucose, glycerol, gelatin, L-alanine, L-aspartic Acid, L-glutamic Acid, pectin, quinic acid, methyl pyruvate, L-lactic acid, citric acid, L-malic acid, tween 40, propionic acid, acetic acid. Strains were resistant to rifamycin SV, lincomycin, vancomycin, tetrazolium violet, tetrazolium blue, lithium chloride. Overall, biochemical characteristics of PPL1, PPL2 and PPL3 strains were in accordance with its close relative *X. sacchari* NCPPB 4341^T^. Biochemical characteristics of *X. albilineans* LMG 494^T^ relative species of *X. sacchari* was taken from literature and included in table 1 [23].

**Table 1:**
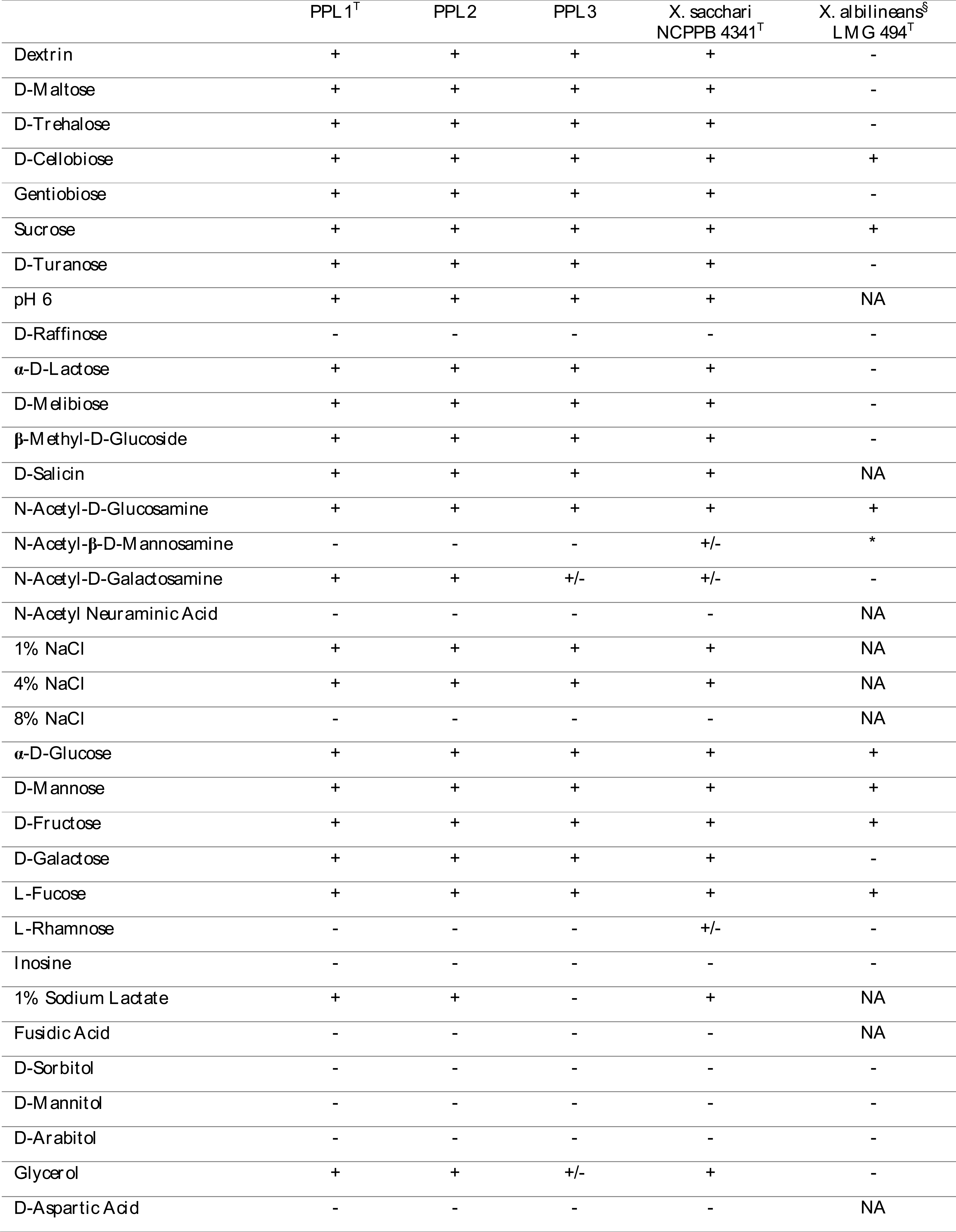
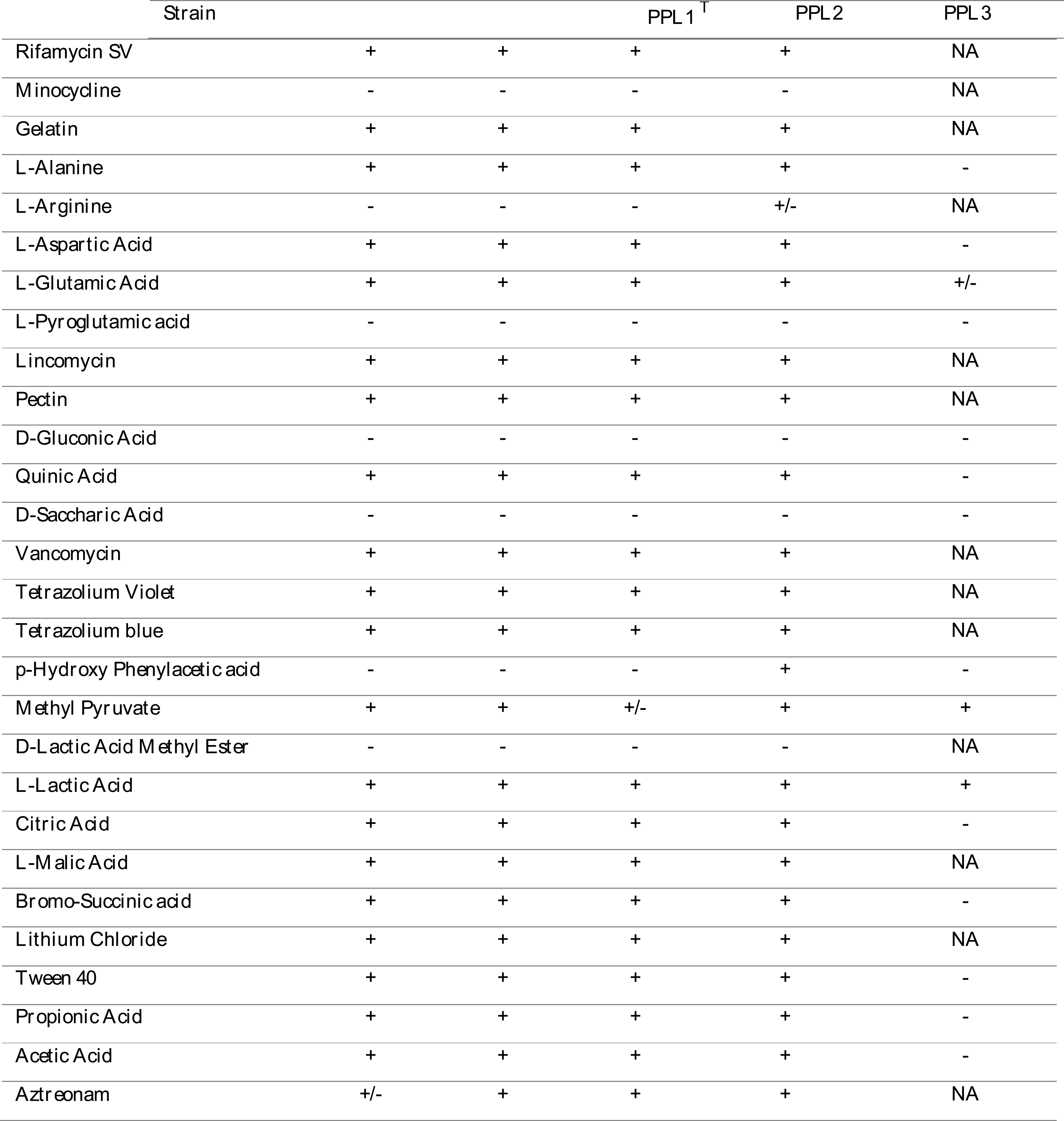
Comparison of biochemical characteristics of PPL1^T^, PPL2, PPL3, and their closest neighbor *X. sacchari* NCPPB 4341^T^. Symbols represents; +: positive, -: negative, +/-: borderline, *§*: *X. albilineans* LMG 494^T^ strain characteristics already reported in [23], NA: data not available in literature.

### Pathogenicity test

Pathogenicity was checked by *in planta* inoculation studies. After 14 days of infection, BXO1 infected leaves showed approx. 13 cm lesion while, PPL1, PPL2, PPL3 and negative control did not show significant infection (approx. 0.5cm lesion) when compared with BXO1 (figure 2). Hence, this clearly reveals that PPL1, PPL2, PPL3 are non-pathogenic to the host.

**Figure 2:**
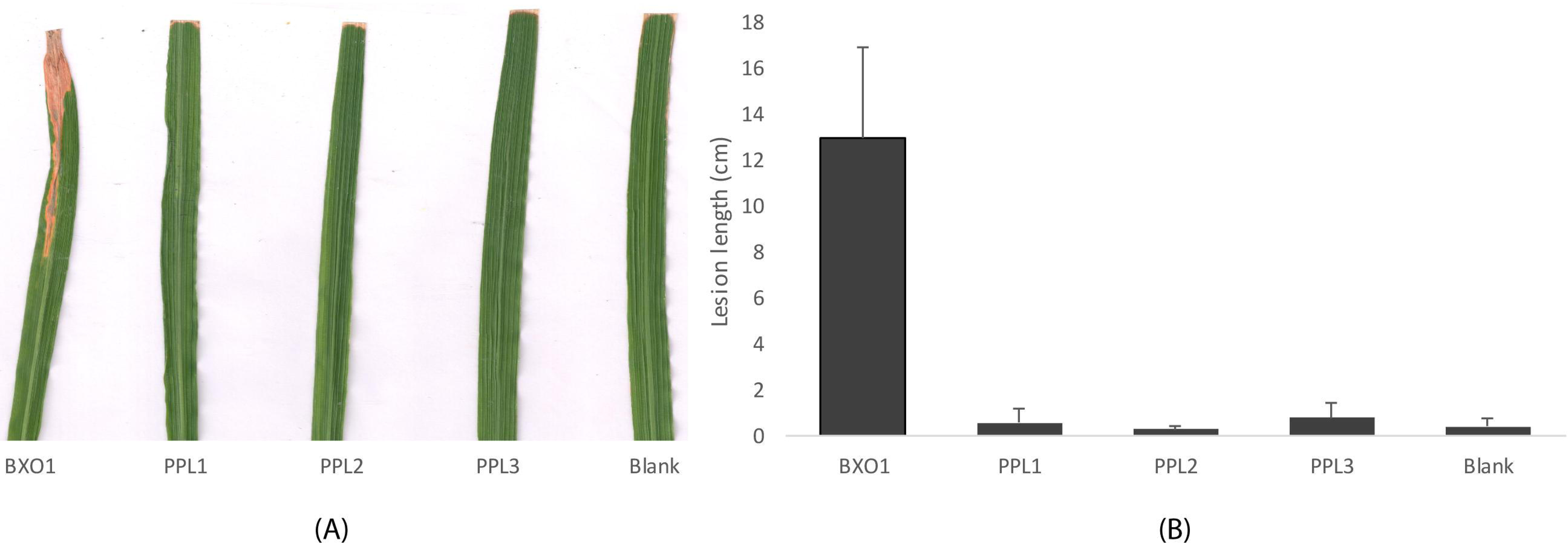
*In planta* infected leaves of rice (Pusa Basmati 1121) (a) leaves showing symptoms of disease after 14 dpi (b) lesion length measured in cm for positive (BXO1), negative (PBS) control and PPL strains. Error bar indicates standard deviation of readings from 10 inoculated leaves and from two independent experiments.

### In-house whole genome sequencing and assembly

Whole genome sequencing of PPL1, PPL2 and PPL3 strains were carried out using in-house Illumina MiSeq platform. The genome size of all strains was approx. 5Mb with genome coverage ranging from 78x to 109x. Assembly statistics for all the strains are given in table 2.

**Table 2:** Genome assembly statistics of PPL1^T^, PPL2 and PPL3 strains.

| S. N. | Strain Name | Completeness/<br>Contamination | Fold (X) | N50 (kb) | Contigs | Genome Size | GC (%) | CDS | rRN A+ tRNA | Isolation source | Accession No. |
| --- | --- | --- | --- | --- | --- | --- | --- | --- | --- | --- | --- |
| 1 | PPL1<sup>T</sup> | 96.31/0.22 | 78 | 32.9 | 332 | 4.8 | 69 | 4149 | 3+51 | Rice seeds | NQYO |
| 2 | PPL2 | 95.79/0.07 | 109 | 48.1 | 200 | 4.9 | 68.8 | 4105 | 2+50 | Rice seeds | NQYP |
| 3 | PPL3 | 98.56/0.55 | 80 | 40.3 | 231 | 4.8 | 68.8 | 4106 | 3+52 | Rice seeds | NMPO |

### Phylogenomic analysis

Core genome tree was constructed using PhyML. For analysis, total 36 species were used (including 32 type and representative strains of different *Xanthomonas* species, along with three strains PPL1, PPL2 and PPL3 isolated in this study). *Stenotrophomonas maltophilia* ATCC13637 was used as an outgroup. Out of 36 species, 27 were clubbed in one group as previously reported [7] including *X. pisi, X. vesicatoria, X. citri, X. codiaei, X. fragariae, X. bromi, X. campestris, X. dyei, X. phaseoli, X. hortorum, X. arboricola, X. cynarae, X. cucurbitae, X. vasicola, X. floridensis, X. perforans, X. euvesicatoria, X. maliensis, X. gardneri, X. axonopodis, X. cassavae, X. nasturtii, X. alfalfae, X. prunicola, X. oryzae, X. melonis*, and *X. populi*. Whereas 8 strains fallen in second group including PPL1, PPL2, PPL3, *X. sacchari, X. theicola, X. translucens, X. hyacinthi, X. albilineans*. PPL1^T^, PPL2 and PPL3 formed a monophyletic clade distinguishing them from other strains. However, *X. sacchari* is the closest neighbour of these strains. Amongst PPL strains, PPL1 and PPL2 are distinct from PPL3 with 100 bootstrap value (figure 3).

**Figure 3:**
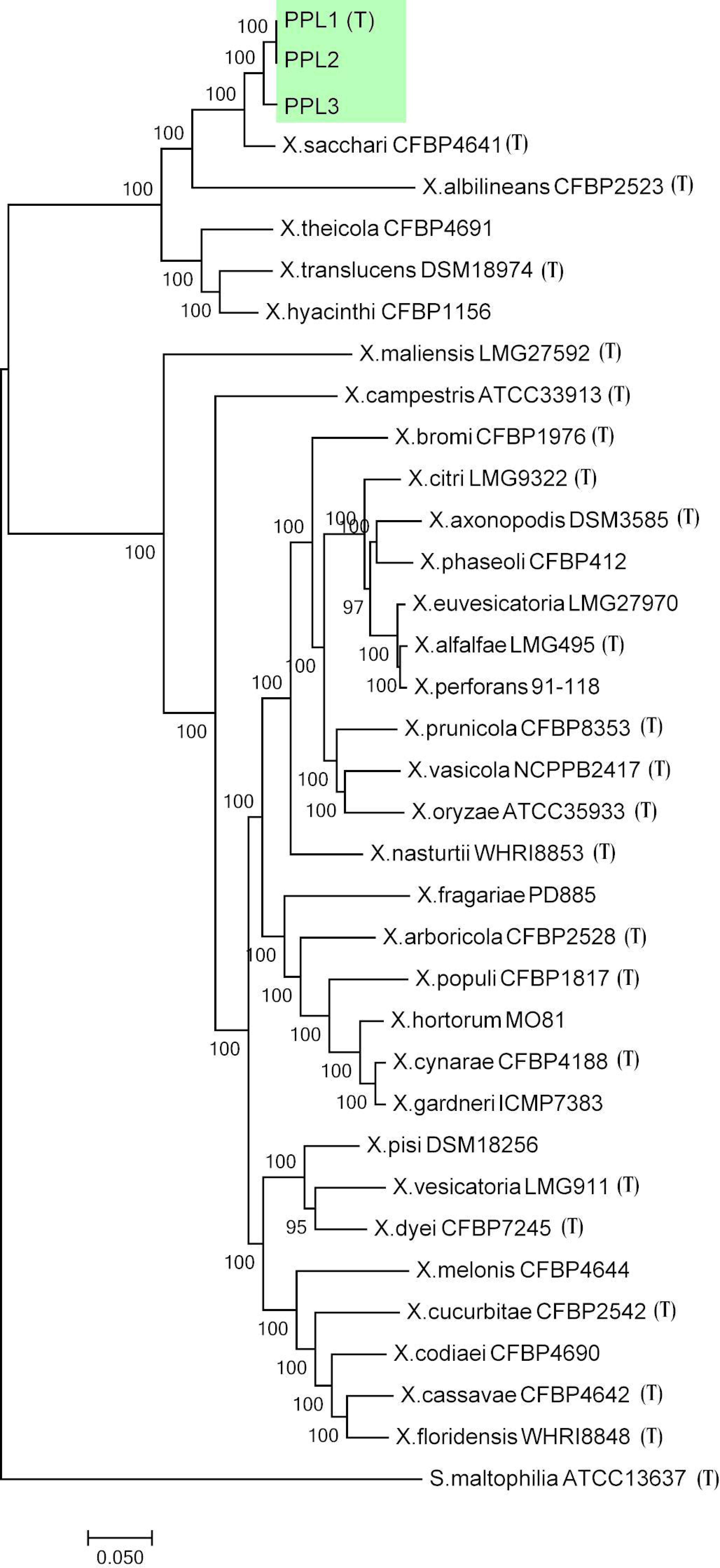
Whole genome based phylogeny considering all type strains and representative strains of genus *Xanthomonas*. The scale bar shows the number of nucleotide substitution per site. PPL1^T^, PPL2 and PPL3 strains (highlighted in coloured box) formed a distinct cluster. *S. maltophilia* ATCC1637 was used as an outgroup.

### Genome based taxonogenomic status

The orthoANI (figure 4) and dDDH values (table 3) of PPL1, PPL2 and PPL3 with type and representative strains of genus *Xanthomonas* species were below the cut-off for species delineation. These strains have *X. sacchari* as their closest relative with ANI values of ∼ 94%, establishing novel species status of these strains. All the PPL1, PPL2 and PPL3 strains showed dDDH values of around 55% with *X. sacchari* and less than 35% with other species of the genus *Xanthomonas*. Interestingly, at the whole genome level, PPL1 and PPL2 strains were found to be clonal (ANI-99.95%, dDDH-99.5%) whereas, PPL3 was not clonal (ANI-97.6%, dDDH-∼78%) however, it belonged to the same species.

**Table 3:**
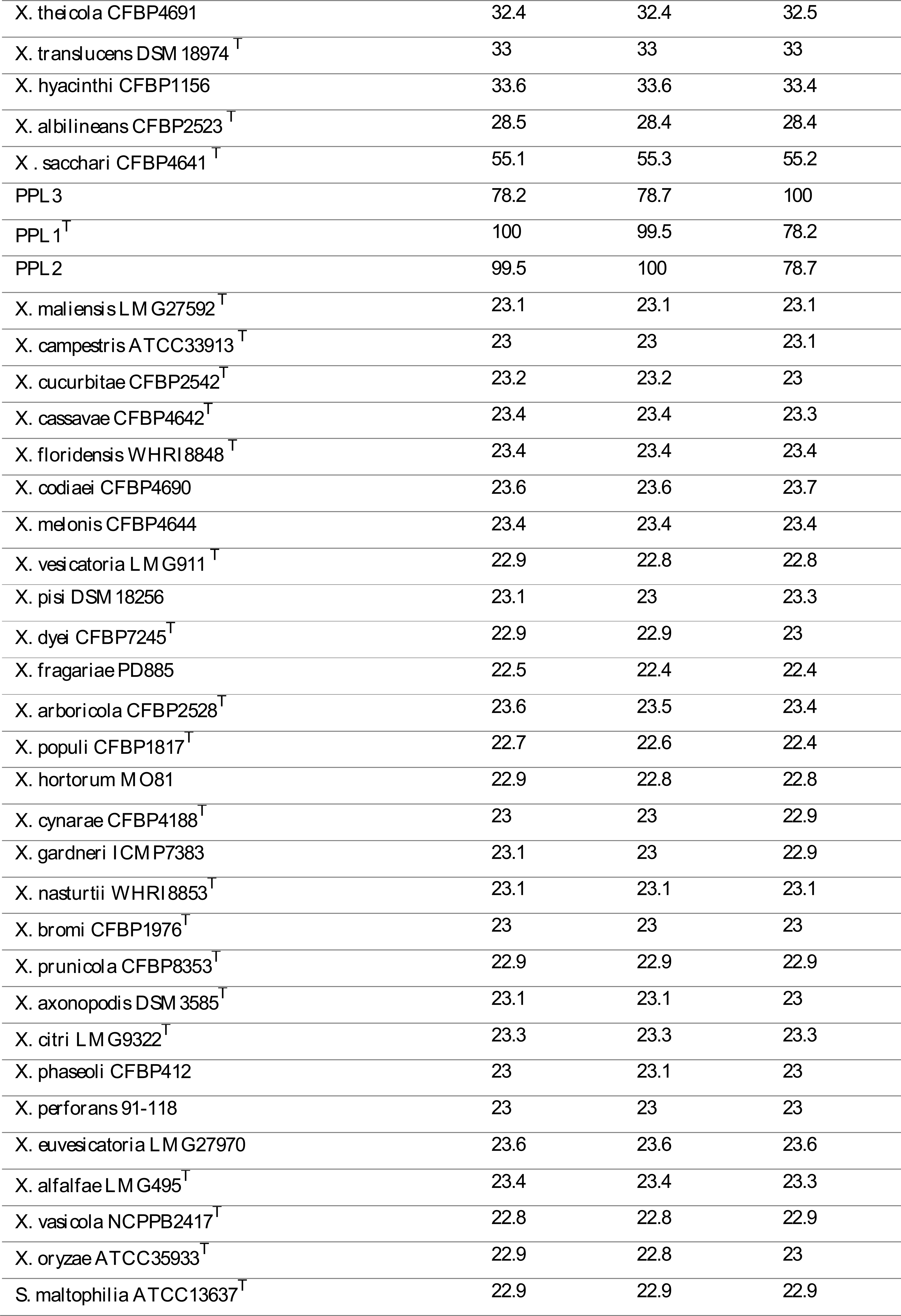
The dDDH values of strains PPL1^T^, PPL2 and PPL3 with other *Xanthomonas* strains.

**Figure 4:**
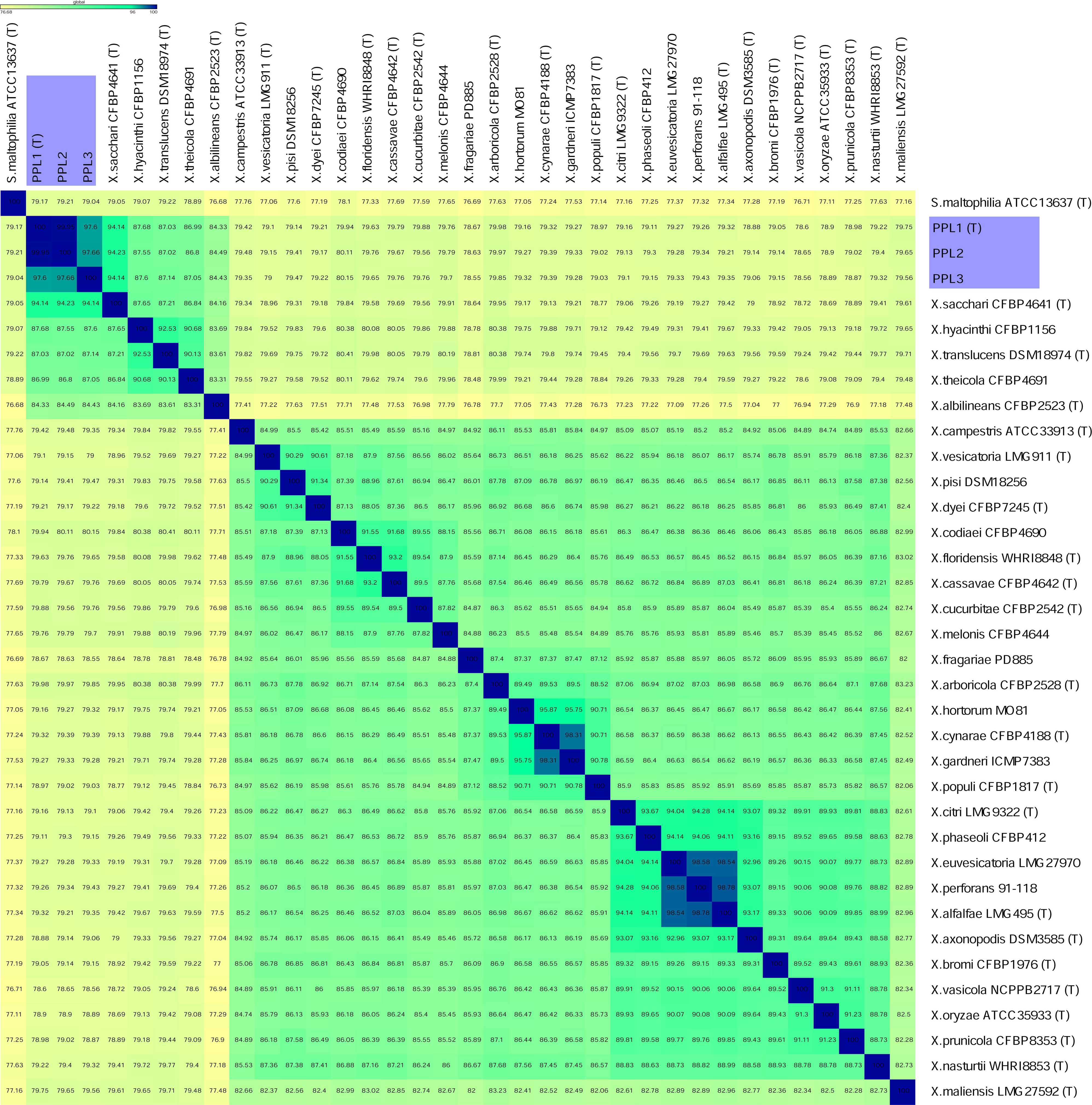
Heat map showing ANI strains values of PPL1^T^, PPL2 and PPL3 with type and representative strains of the genus *Xanthomonas*.

### Description of *Xanthomonas sontii* sp. nov

*Xanthomonas sontii* (N.L. masc. gen. n. *sontii* named in honour of Ramesh V. Sonti, a renowned Indian bacterial and plant molecular geneticist).

Cells are Gram-negative, aerobic, rod shape, motile and form yellow, round, smooth, convex and circular colonies after 24hrs. Cells can grow on nutrient agar (NA), peptone sucrose agar (PSA) and glucose yeast extract calcium carbonate agar (GYCA) media. Optimum temperature for growth is 28 °C. They are able to utilize of D-maltose, D-trehalose, D-cellobiose, gentiobiose, sucrose, D-turanose, α-D-lactose, D-melibiose, β-methyl-D-glucoside, D-salicin, N-acetyl-D-glucosamine, α-D-glucose, D-mannose, D-fructose, D-galactose, L-fucose, glycerol, gelatin, L-alanine, L-aspartic acid, L-glutamic acid, pectin, quinic acid, methyl pyruvate, L-lactic acid, citric acid, L-malic acid, tween 40, propionic acid, acetic acid. Strains were able to grow at pH 6.0 and resistant to 1% NaCl, 1% sodium lactate, and antibiotics like rifamycin SV, lincomycin, vancomycin, tetrazolium violet, tetrazolium blue, lithium chloride. Taxonogenomic and phylogenomic analysis revealed distinctness of these species with orthoANI and dDDH values below established cutoff values i.e. 96% for ANI and 70% for dDDH. Core genome tree analysis showed separate grouping PPL1, PPL2 and PPL3 from other *Xanthomonas* strains. Further, amongst PPL strains PPL3 differ at clone level forming distinct clade than PPL1 and PPL2. Therefore, we propose PPL1, PPL2 and PPL3 as novel species *X. sontii* of the genus *Xanthomonas* with PPL1 as type strain PPL1^T^ (CFBP8688^T^ = ICMP23426^T^ = MTCC12491^T^).

## Acknowledgements

We thank Randeep sharma for providing technical assistance with electron microscopy. All the work in present study was supported by “(MICRA) – Mega-genomic insights into co-evolution of rice and its Microbiome” (MLP0020).

## Abbreviations

OrthoANI: Orthologous average nucleotide identity
dDDH: digtal DNA-DNA hybridization
NA: nutrient agar
PSA: peptone sucrose agar
GYCA: glucose yeast extract calcium carbonate agar
PBS: phosphate buffer saline
TSBA: tryptic soy agar
MCS: MiSeq control software

